# Gender-specific MetS prediction using pathophysiological determinants: Beyond diagnostic constraints

**DOI:** 10.1101/2025.06.10.658812

**Authors:** Karishma Desai, Indra Kundu, Vivek Verma, M S Aravind, Manu Sudhakar, Jaideep Menon, Susan Idicula-Thomas

## Abstract

Metabolic syndrome (MetS), constituted by obesity, hyperglycemia, hypertension, and dyslipidemia, is a growing public health concern due to its associated risk with cardiovascular and other metabolic disorders. Early-stage detection and prevention of MetS are key factors in an effective management strategy. Thus, the current study aimed to develop a risk score for MetS progression based on features pertaining to its pathophysiology, that can facilitate early detection and treatment strategies. A multivariate logistic regression model was developed using a representative feature from MetS pathophysiological pathways of inflammation, endothelial dysfunction, and hepatic dysregulation. The model was built on NHANES dataset and validated in Chinese and Indian datasets. The model performance evaluated using ROC resulted in an AUC of 0.81 in training and 0.87 and 0.79 for Chinese and Indian validation datasets, respectively. In conclusion, a MetS predictive model built on three MetS pathophysiological determinants was developed and validated in distinct datasets.

**Highlights:**

1. A MetS prediction model, trained on NHANES data, is validated with two datasets of varied ethnicity.
2. The three parameter model, independent of MetS diagnostic criteria, is reflective of inflammation, endothelial dysfunction, and hepatic dysregulation.
3. The model achieved an AUC of 0.81 in NHANES data; 0.87 and 0.79 for Chinese and Indian validation datasets respectively.

## Introduction

Metabolic syndrome (MetS) is characterized by presence of central obesity, hyperglycemia, hypertension, increased triglycerides, and decreased high density lipoprotein (HDL). The global prevalence of MetS ranges from 14.8% to 46.4% depending on the definition used for diagnosis and the region studied^1^ and the prevalence of MetS has increased over time across diverse geographical locations^2–4^. MetS is associated with increased risk of co-morbidities including type 2 diabetes, cardiovascular diseases, non-alcoholic fatty liver disease, and cancers ^5^. Early stratification of individuals, based on their risk of developing MetS, will enable timely intervention, preventing its onset and progression to associated comorbidities. Several machine learning (ML)-based prediction models have been developed for MetS using features such as basic demographic details, anthropometric measurements, biochemical data, and genetic factors^6–8^. These models have incorporated MetS diagnostic parameters such as waist circumference, HDL, triglycerides in their training and therefore are not truly predictive in nature and lack representation of MetS progression.

Thus, the objective of the current study was to develop a risk score for MetS progression based on features pertaining to its pathophysiology, that can facilitate early detection and treatment strategies. The model trained on National Health and Nutrition Examination Survey (NHANES) was validated in two distinct populations groups of Chinese and Indian origin.

## Methods

Population survey-based datasets from 1999 to 2020 (N = 1,16,876) were downloaded from NHANES webpage (https://wwwn.cdc.gov/nchs/nhanes/). The dataset was restricted to individuals aged 31–55 years to exclude an overly healthy younger control group and ensure a more clinically relevant comparison population; missing data for age, clinical and biochemical features that are associated with MetS diagnosis and presence of co-morbidities, or previous history of hypertension, cholesterol, or diabetes were removed (Figure S1). This subset of the dataset was used to develop a model that can classify individuals as control or MetS based on blood-based biochemical features (excluding the diagnostic parameters).

### a) Feature selection

MetS diagnostic parameters, such as waist circumference, blood pressure, fasting blood sugar, triglyceride and HDL levels, were removed. In accordance with the well-characterized pathophysiology of MetS, wherein inflammation, hepatic dysregulation, and endothelial dysfunction are established as central mechanisms, one representative feature was selected from each pathway to build a prediction model.

### b) Model generation and validation

A multivariate logistic regression model with three unique features differentiating control vs MetS was built. The NHANES dataset was used to evaluate the model performance and two independent datasets from China 9 and India (Amrita hospital, Kerala) were used for assessing the model’s reproducibility and generalizability. The Indian validation dataset was from a prospective cross-sectional study design, comprised of data collected from individuals visiting Amrita Institute of Medical Sciences (AIMS), Kerala for a comprehensive health check-up (supplementary methods). The study design was approved by Institutional ethics committee of AIMS, Kerala (ECASM-AIMS-2022-173) and ICMR-NIRRCH (project number: 476/2022). The logistic regression equation was subjected to gender-specific ROC analysis such that the value associated with the maximum sum of sensitivity and specificity (i.e., maximising the Youden index) was selected as the “optimal” threshold value for NHANES dataset.

## Results

### a) Model displayed AUC >0.8 for training and validation datasets

Of the 7,666 participants in the NHANES dataset, 2,345 (30.59%) and 610 (7.95%) were assigned to the MetS and control groups respectively. WBC, UA, and ALT/AST parameters were used to build a multivariate logistic regression model. The predictive accuracy of these parameters individually and in combination was evaluated and highest accuracy was attained with three parameter model (Figure S2).

The model trained on NHANES dataset achieved an AUC of 0.81 in differentiating between MetS cases and control datasets. The model validation in Chinese and Indian populations resulted in AUC of 0.87 and 0.82 respectively (Figure S3). The equation generated by multivariate logistic regression model was logit(Y) = 0.30*WBC + 0.007*UA + 2.86*(ALT/AST) where WBC is expressed as 10^9^/L and UA as μmol/L.

### b) Gender specific threshold for MetS prediction

The logistic regression model generated gender-specific thresholds of risk score as 7.47 (AUC=0.83) for men and 6.35 (AUC=0.85) for women from the NHANES dataset. The threshold validation in Chinese dataset generated an AUC of 0.81 for males and 0.72 in females. In the second validation dataset of Indian origin, the risk score threshold resulted in AUC of 0.81 in males and 0.70 in females (Figure 1). Additionally, the threshold of risk score was evaluated in the pre-MetS conditions (one and two component states) and the distribution is shown in Figure 2. The risk score progressively increased from control to pre-MetS to MetS states across genders and ethnicities. For males, the threshold of 7.47 seemed to be effective and consistent across all ethnic groups examined, highlighting its utility in tracking disease progression. In females, while the trend of risk score proportionately increasing with disease progression remained consistent, the threshold seemed to be higher in Chinese and Indian dataset as compared to NHANES, underscoring the importance of population-specific derivation of risk score threshold or cut-off values.

**Figure 1:**
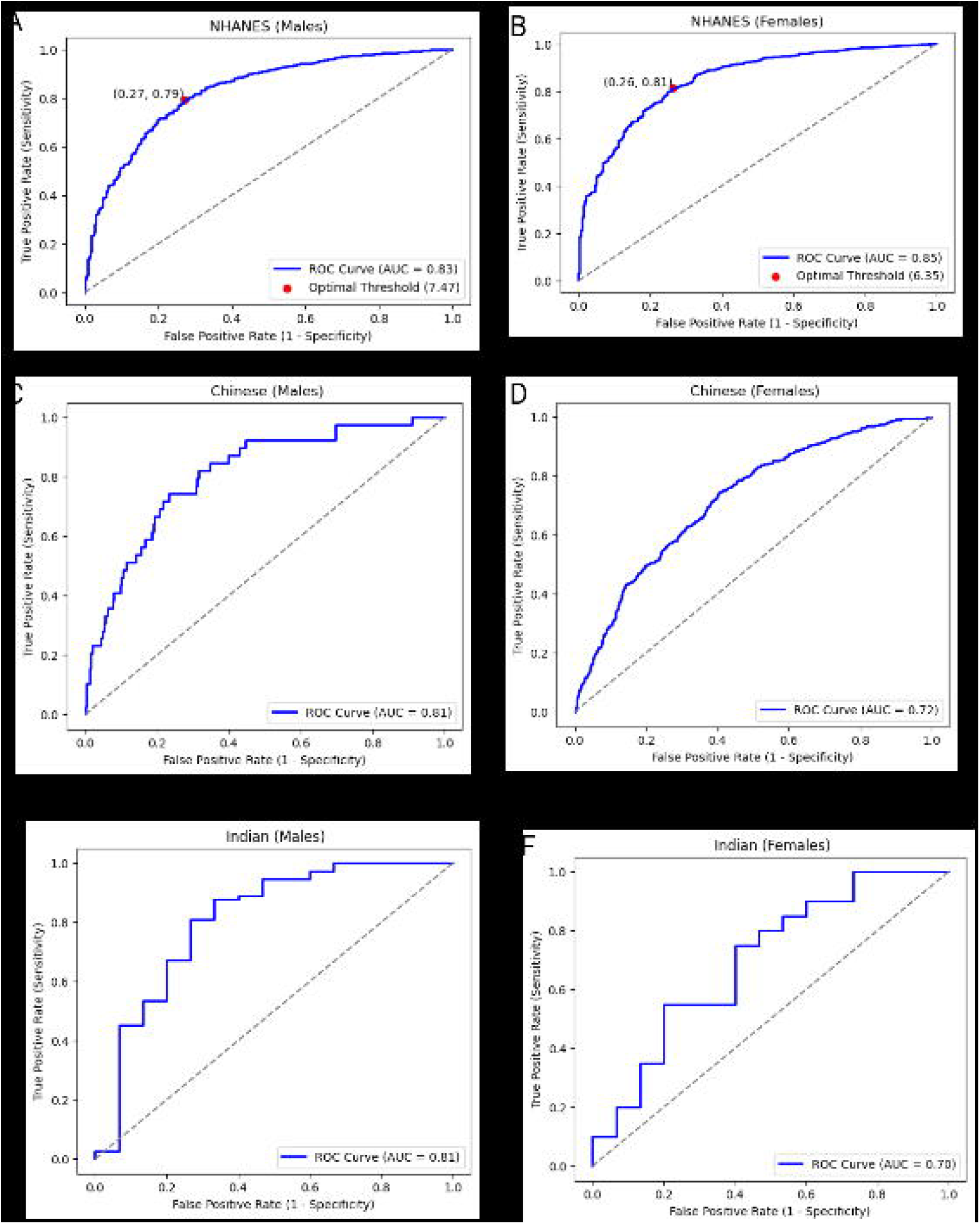
Model and risk score performance in training and validation datasets. ROC analysis for model performance in (A) males and (B) females of the NHANES dataset with optimal threshold defined by Youden index. ROC analysis for evaluating risk score performance in (C) Chinese males, (D) Chinese females, (E) Indian males, and (F) Indian females.

**Figure 2:**
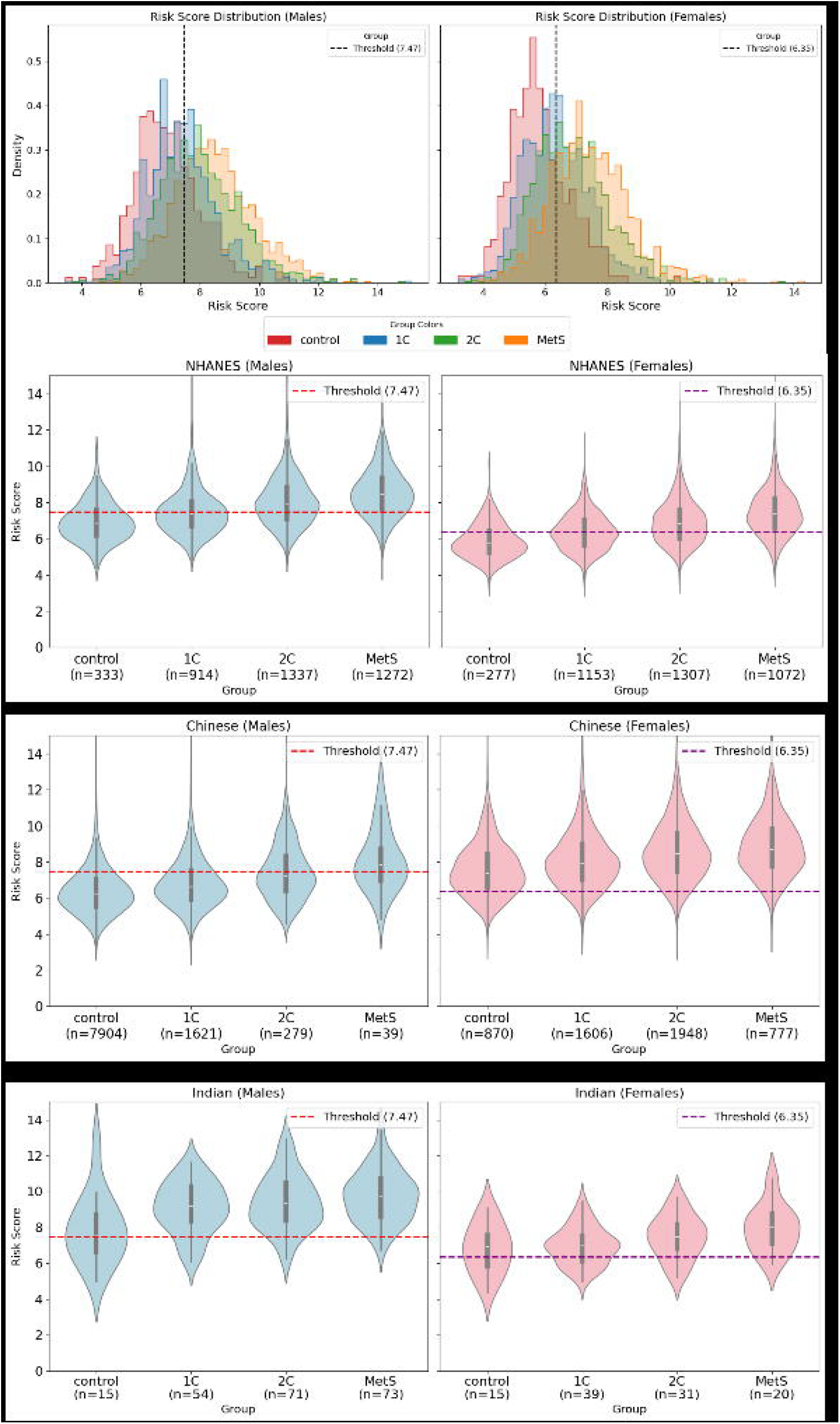
Gender-wise distribution based on risk score. (A)A density distribution plot for the three datasets based on their phenotype (group) for males and females. A box and violin plot for risk score against the phenotypes for (B) NHANES, (C) Chinese, and (D) Indian datasets. (B-D) The left panel depicts distribution for males and right depicts for females respectively.

## Discussion

The rising prevalence of MetS, its components, and associated comorbidities contribute significantly to the burden of non-communicable diseases. Developing and validating a clinical risk prediction model for MetS can aid healthcare professionals in early disease diagnoses, identifying high-risk individuals, and enabling timely intervention for better patient outcomes. This study developed a gender-specific MetS risk score using a multivariate logistic regression model. The model, which did not rely on MetS-defining parameters, could effectively differentiate between control and MetS in three independent datasets. The model and risk score thresholds were trained on the NHANES dataset, and validated using distinct Chinese and Indian population datasets.

The multivariate model was developed with three pathophysiological features - WBC, UA, and ALT/AST ratio. The model achieved an AUC of >0.8 in training and validation datasets. Despite being trained on individuals aged 31–55 years, the model achieved a comparable AUC on the 18–50 years Indian validation set, suggesting its potential applicability to younger age groups. Various research groups have generated MetS prediction models utilizing non-invasive parameters like BMI, waist-to-hip ratio, blood pressure readings, and questionnaire-based data^10,11^. However, as these features are linked to the diagnostic criteria for MetS, such models primarily capture ongoing metabolic changes rather than acting as early indicators of risk. Notably, while these models achieved a comparable AUC with a larger set of features and defining parameters (n = 5-20), the present study demonstrated equivalent performance using only three features, none of which are associated with MetS diagnosis. It is reassuring that the predictive features of the model reflected three key aspects of MetS pathophysiology: systemic inflammation (WBC), endothelial dysfunction (UA) and hepatic dysregulation (ALT/AST). It is noteworthy that the three parameters – WBC, UA and AST/ALT tests are routinely included in health check-up packages, making them affordable as compared to biomarkers such as hsCRP that are reported for systemic inflammation and MetS^12^. Monitoring the disruption in these mechanisms can facilitate early prediction of MetS, as evidenced by the distribution of risk scores across the disease states. The score exhibited a progressive increase corresponding to the accumulation of components, beginning from a healthy state with no components to a MetS state with three or four components. (Figure 2).

Since UA levels are naturally lower in females compared to males, gender-specific thresholds for the risk score were established using the NHANES training dataset. These thresholds performed well for males across validation datasets. However, in females, although the risk score rose proportionately in MetS cases as expected, the threshold seemed to require adjustment in the validation datasets, as the control group displayed higher than expected scores (Figure 2). Mann-Whitney tests comparing WBC, UA, and ALT/AST levels among control groups across the three datasets revealed significant differences (Figure S4), highlighting the necessity of developing population-specific thresholds. Some of the observed variation may be due to differences in ethnicity, lifestyle, or genetic background, which were not evaluated in the current analysis. Further research is required to address these aspects.

We recognize the importance of validating the risk score using a larger dataset to enhance its robustness and broader applicability. Furthermore, validation through cohort studies is critical to assess its effectiveness in predicting the progression from pre-MetS states to MetS. These efforts are critical for demonstrating the application of the prediction model in diverse populations, enabling early interventions and lifestyle modifications to prevent the advancement to MetS.

## Authors contribution

Karishma Desai: Writing – review & editing, Writing – original draft, Investigation, Formal analysis. Indra Kundu: Writing-review & editing, Methodology, Investigation, Formal analysis. Vivek Verma: Writing – review & editing, Methodology, Formal analysis. Aravind MS: Writing – review & editing, Methodology. Manu Sudhakar: Writing – review & editing, Methodology. Jaideep Menon: Writing – review & editing, Methodology. Susan Idicula-Thomas: Writing – review & editing, Supervision, Resources, Project administration, Funding acquisition, Formal analysis, Conceptualization.

## Acknowledgements

The authors would like to acknowledge all participants for their participation in the study.

## Data availability statement

The data used in the study were derived from the National Health and Nutrition Examination Survey (NHANES) accessed through the NHANES website at https://www.cdc.gov/nchs/nhanes/index.htm. The Chinese dataset was accessed through the DOI https://doi.org/10.1186/s12902-022-01121-4. All data supporting the findings of this study are available from the corresponding author upon reasonable request.

## References

1. Noubiap JJ, Nansseu JR, Lontchi-Yimagou E, et al. Geographic distribution of metabolic syndrome and its components in the general adult population: A meta-analysis of global data from 28 million individuals. Diabetes Res Clin Pract. 2022;188. doi:10.1016/J.DIABRES.2022.109924

2. Yang C, Jia X, Wang Y, et al. Trends and influence factors in the prevalence, intervention, and control of metabolic syndrome among US adults, 1999-2018. BMC Geriatr. 2022;22(1). doi:10.1186/S12877-022-03672-6

3. Park D, Shin MJ, Després JP, Eckel RH, Tuomilehto J, Lim S. 20-Year Trends in Metabolic Syndrome Among Korean Adults From 2001 to 2020. JACC Asia. 2023;3(3):491–502. doi:10.1016/J.JACASI.2023.02.007

4. 1. Feng T, Zheng J, Wang X, et al. Decadal Trends in the Prevalence of Metabolic Syndrome in Economically Developed Regions in China. J Endocr Soc. 2024;8(8). doi:10.1210/JENDSO/BVAE128

5. Pedro-Botet J, Ascaso JF, Barrios V, et al. COSMIC project: consensus on the objectives of the metabolic syndrome in clinic. Diabetes Metab Syndr Obes. 2018;11:683–697. doi:10.2147/DMSO.S165740

6. Shin D. Prediction of metabolic syndrome using machine learning approaches based on genetic and nutritional factors: a 14-year prospective-based cohort study. BMC Med Genomics. 2024;17(1). doi:10.1186/S12920-024-01998-1

7. Shin H, Shim S, Oh S. Machine learning-based predictive model for prevention of metabolic syndrome. PLoS One. 2023;18(6). doi:10.1371/JOURNAL.PONE.0286635

8. Zou G, Zhong Q, OUYang P, Li X, Lai X, Zhang H. Predictive analysis of metabolic syndrome based on 5-years continuous physical examination data. Sci Rep. 2023;13(1). doi:10.1038/S41598-023-35604-8

9. Zhang Y, Zhang X, Razbek J, et al. Opening the black box: interpretable machine learning for predictor finding of metabolic syndrome. BMC Endocr Disord. 2022;22(1). doi:10.1186/S12902-022-01121-4

10. Sherman-Hahn S, Izkhakov E, Perlman S, Ziv-Baran T. A new metabolic syndrome prediction model for self-evaluation as a primary screening tool in an apparently MetS-free population. Prev Med (Baltim). 2023;175. doi:10.1016/J.YPMED.2023.107701

11. Kim J, Mun S, Lee S, Jeong K, Baek Y. Prediction of metabolic and pre-metabolic syndromes using machine learning models with anthropometric, lifestyle, and biochemical factors from a middle-aged population in Korea. BMC Public Health. 2022;22(1). doi:10.1186/S12889-022-13131-X

12. Mirhafez SR, Ebrahimi M, Saberi Karimian M, et al. Serum high-sensitivity C-reactive protein as a biomarker in patients with metabolic syndrome: evidence-based study with 7284 subjects. Eur J Clin Nutr. 2016;70(11):1298–1304. doi:10.1038/EJCN.2016.111

