## Supplementary methods for "Gender-specific MetS prediction using pathophysiological determinants: Beyond diagnostic constraints"

**Supplementary section:**

1. External validation datasets
i. *Dataset from Zhang Y et.al., 2022 (Chinese dataset)*

The first validation dataset was downloaded from the supplementary section of Zhang Y et.al., 2022^1^. This dataset was obtained from chain of health screening institutions in Urumqi, Xinjiang Uygur Autonomous Region, China. The data was collected between 2017-2019. The data contains basic demographic details, questionnaire surveys, and physical and laboratory investigations of 39,134 individuals. The authors have provided the dataset open access, in the supplementary section which was downloaded for further analysis. The dataset was filtered for age 31-55 years (24,919 of 39,134 individuals) and individuals with previous history of liver diseases, hypertension, or diabetes were excluded for further analysis. 15,070 (9,865 males and 5,205 females) individuals were included in the analysis.

ii. *Cross-sectional data from a tertiary care hospital in India (Indian dataset)*

The second external validation set comprised of data collected from individuals attending the outpatient departments of Amrita Institute of Medical Sciences (AIMS), Kerala. Individuals were recruited if they were in the age group of 18-50 years and were either healthy or treatment-naïve and diagnosed with MetS or any one or combination of its components as per the modified Harmonized definition. Individuals with any other co-morbidities such as liver diseases, type II diabetes, thyroid, diabetes or CVDs were excluded during the screening. A total of 300 individuals were recruited and included in the validation set. Blood samples under fasting conditions were collected from individuals and were used for blood based biochemical assessments.

The study design was approved by Institutional ethics committee of AIMS, Kerala (ECASM-AIMS-2022-173) and ICMR-NIRRCH (project number: 476/2022).

2. **Definitions**:

Individuals were assigned to subgroups of MetS and its components (Obs, Hyg, Hyp, and Dys or their combinations) based on the modified Harmonized definition^2^.

a) Obs: Waist circumference (WC): >90 cm for men and >80 cm for women
b) Hyg: Fasting blood glucose: >100mg/dL or >5.6mmol/L
c) Hyp: Blood pressure >140/90 mmHg
d) Dys: Triglycerides >150mg/dL or >1.7 mmol/L and/or HDLc <40 mg/dL or 1.03 mmol/L for men, <50 mg/dL or 1.29 mmol/L for women

Individuals with levels for all four parameters below the threshold as mentioned above were assigned to *control* groups. Individuals with one or two components were categorized as *pre-MetS* and individuals meeting 3 or 4 criteria were grouped into *MetS*.

**Supplementary figures**:

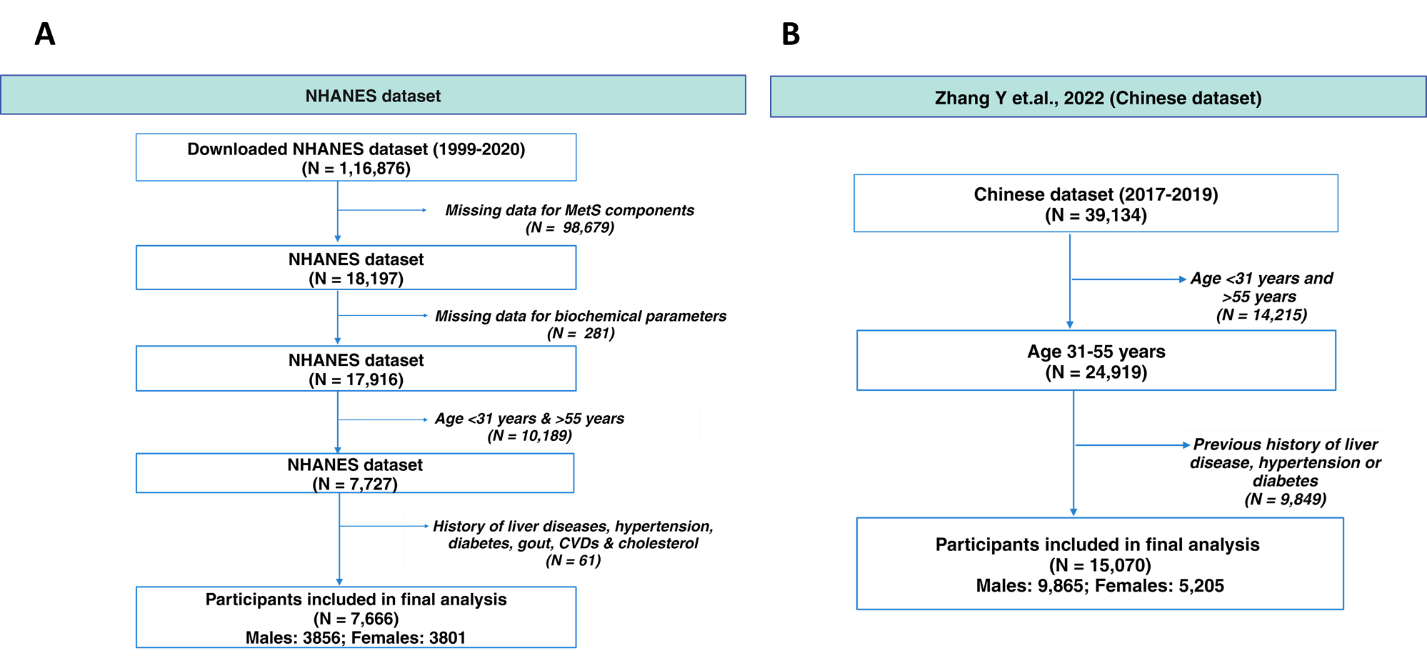

**Figure S1: Flowchart of participant selection from the datasets.**

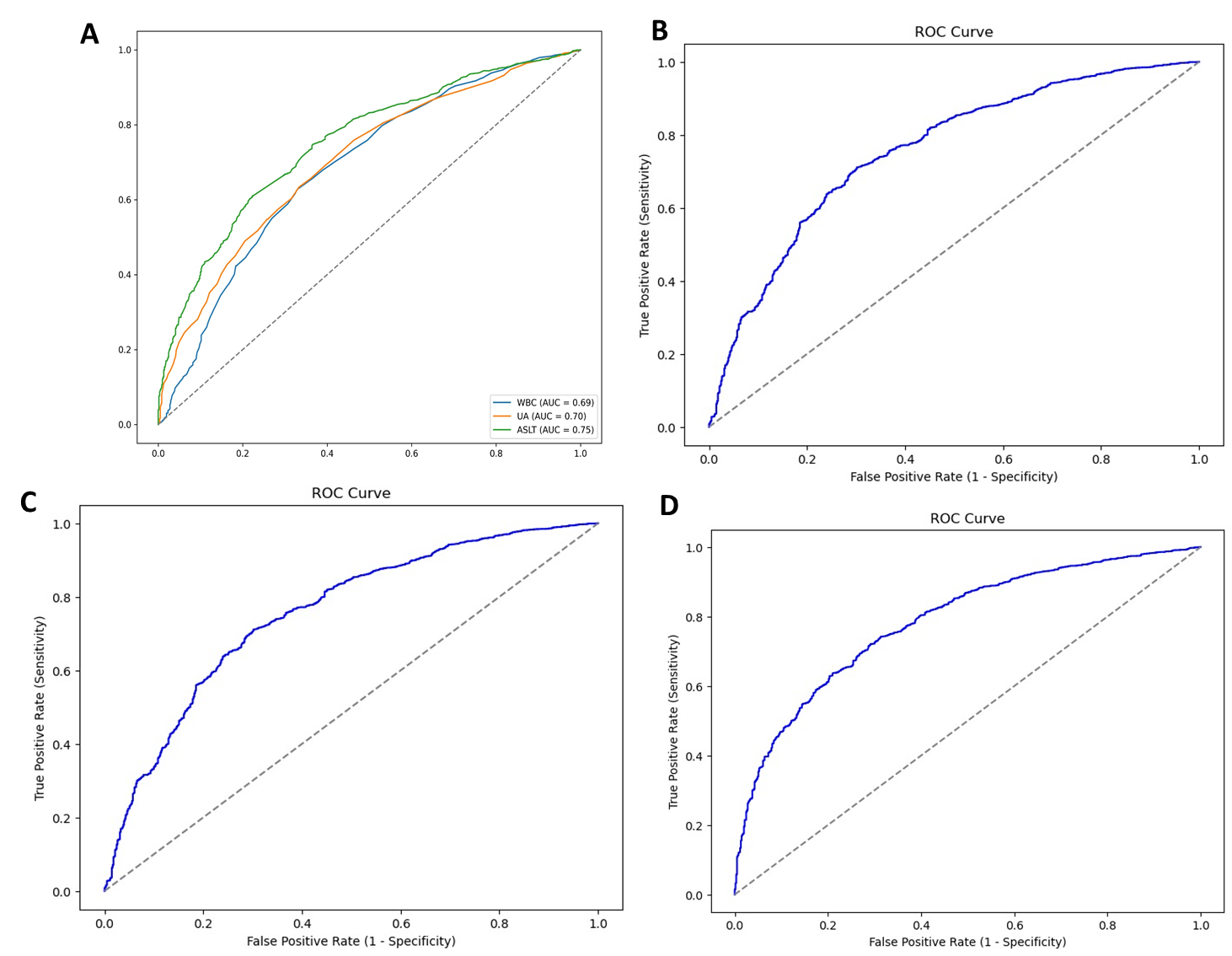

**Figure S2: ROC curves for univariate and bivariate logistic regression models of MetS pathophysiological parameters**. A) Univariate AUC of model features B) AUC of 0.76 for WBC & UA C) AUC of 0.78 for WBC & ALT/AST, D) AUC of 0.78 for UA & ALT/AST

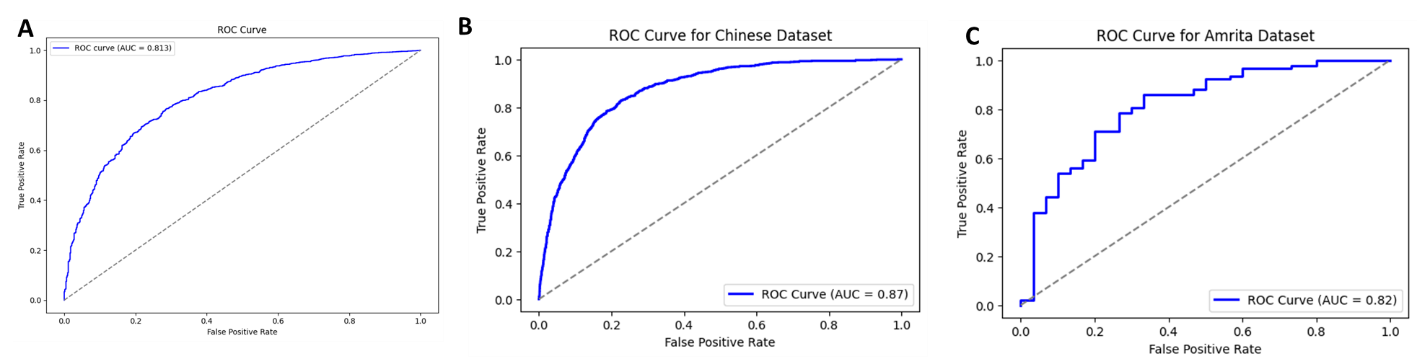

**Figure S3: ROC curves for multivariate logistic regression model built on WBC, UA, ALT/AST.** ROC curve with corresponding AUC values for A) NHANES dataset (AUC = 0.81) B) Chinese dataset (AUC = 0.87) and C) Indian dataset (AUC = 0.82)

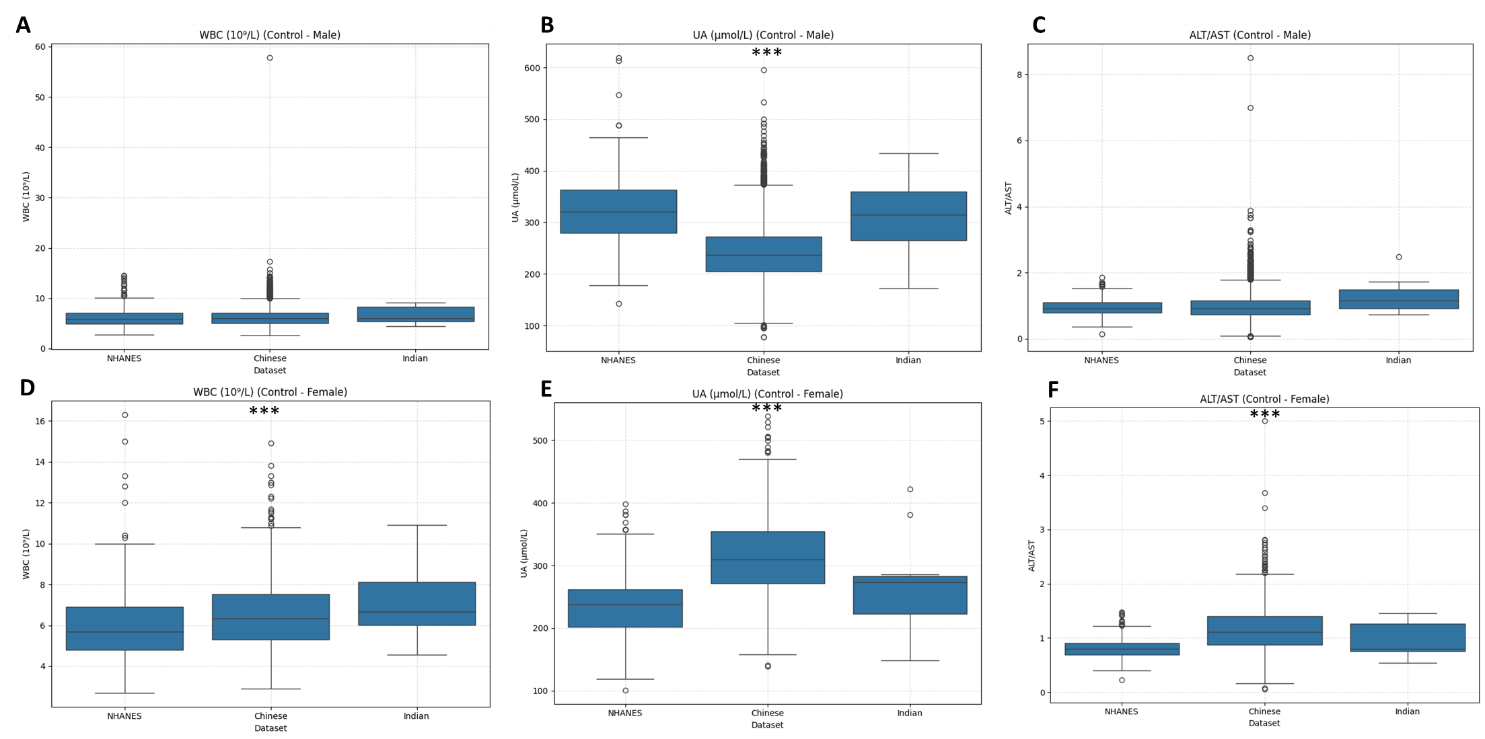

**Figure 4: Box and whiskers plot depicting the distribution of model parameters in the control group of the three datasets.** Gender specific distribution of model parameters in males (A-C) and females (D-F) for WBC (10^9^/L) (A and D), UA (µmol/L) (B and E), ALT/AST (C and F). Statistically significant is considered at p<0.001 compared to NHANES dataset (training).

**References**:

1. Zhang Y, Zhang X, Razbek J, et al. Opening the black box: interpretable machine learning for predictor finding of metabolic syndrome. BMC Endocr Disord. 2022;22(1):214. Published 2022 Aug 26. doi:10.1186/s12902-022-01121-4
2. Alberti KG, Eckel RH, Grundy SM, et al. Harmonizing the metabolic syndrome: a joint interim statement of the International Diabetes Federation Task Force on Epidemiology and Prevention; National Heart, Lung, and Blood Institute; American Heart Association; World Heart Federation; International Atherosclerosis Society; and International Association for the Study of Obesity. Circulation. 2009;120(16):1640-1645. doi:10.1161/CIRCULATIONAHA.109.192644
